# Activation of complement protein C3 depicts the dynamic tissue damage of the cochlea after noise trauma

**DOI:** 10.1101/2025.04.20.649717

**Authors:** Zixu Guo, Benjamin J Seicol, Mina Shenouda, Angela Wang, Katy Garrity, Shengyin Lin, Ruili Xie

## Abstract

Hearing loss is characterized by dysfunction and pathological changes of the cochlea, including the loss of sensory hair cells and connected synapses. Growing evidence has revealed that the immune system plays critical roles in shaping cochlear responses to tissue injury and the clearance of damaged cells under pathological conditions such as noise-induced hearing loss (NIHL). However, the underlying mechanisms that regulate this process remain unclear. As a crucial part of the innate immune system, the complement system is widely known to respond to tissue damage and help remove cellular debris. Such functions are yet to be fully explored in the auditory system, especially in the cochlea. Among various complement factors, the activation of complement protein C3 is an essential functional hub of the complement cascade, resulting in the phagocytosis of dead or dying cells. In this study, we sought to investigate whether C3 exists in the cochlea and what roles it might play upon noise injury. To test this, we studied C3 activation using immunohistochemistry in the cochleae of various mouse models that were noise exposed to an octave band noise (8–16 kHz) for two hours at 112 dB SPL. Mice were sacrificed at various post exposure times to examine cochlear pathology. We found damage dependent C3 activity in the organ of Corti in mice after noise exposure (NE), especially in damaged outer hair cell (OHC) area. Specifically, C3 opsonized damaged cellular structures include degenerating OHCs, OHC debris, ribbon synapses, and efferent synaptic boutons, and the temporal dynamics of the opsonization closely resemble the timing of cochlear tissue damages post exposure. Unexpectedly, C3-deficient mice exhibited similar ABR thresholds and OHC counts at baseline as wild type mice, as well as comparable ABR threshold shifts and OHC loss after NE. However, we observed significantly reduced MOC terminal bouton degeneration in C3-deficient mice upon 112 dB SPL noise injury, indicating that C3 does modulate tissue damage and clearance, at least at the MOC terminals. Our results suggest that complement C3 is dynamically activated in the cochlea upon noise injury, which may play significant roles in damage recognition and further immune recruitment during the development of NIHL.

## Introduction

Hearing loss is one of the most prevalent sensory deficits largely due to pathological changes and dysfunction of the cochlea. A range of insults to the cochlea—such as excessive noise exposure (NE)—can lead to pathological outcomes, including shifts in hearing thresholds (Kujawa & Liberman, 2009, 2015), sensory hair cell death (Abrashkin et al., 2006; Ping Yang et al., 2004), cochlear synaptopathy (Kujawa & Liberman, 2009, 2015), degeneration of the medial olivocochlear (MOC) efferent system (Canlon et al., 1999; Fuchs & Lauer, 2019), and broader pathological changes in cochlear tissue (Wang et al., 2002). In recent years, growing evidence has revealed that the cochlear immune system plays a critical role in shaping cochlear response to tissue injury and the clearance of damaged cells under pathological conditions such as noise-induced hearing loss (NIHL). NE triggers the release of pro-inflammatory mediators, induces the infiltration of circulating immune cells, and activates resident cochlear macrophages (Fujioka et al., 2006; Manickam et al., 2023, 2024; Pan et al., 2024; Rai et al., 2020; Wood & Zuo, 2017; Yang et al., 2015). However, the specific molecular pathways that govern damage recognition, debris clearance, and the downstream effects of cochlear immune activation remain poorly understood.

The complement system is a strong candidate for mediating these effects. It is a key component of the innate immune response and consists of a cascade of soluble proteins that coordinate immune surveillance and clearance of damaged tissue (Dunkelberger & Song, 2010; Fishelson et al., 2001; Ricklin et al., 2016; Sarma & Ward, 2011; Vandendriessche et al., 2021). Upon activation, the complement cascade mediates opsonization and phagocytosis of cellular debris, and coordinate inflammatory signaling (Dunkelberger & Song, 2010; Fishelson et al., 2001; Ricklin et al., 2016; Sarma & Ward, 2011; Vandendriessche et al., 2021). Beyond its classical immune functions, the complement system is also active in the nervous system, where it participates in synaptic pruning and circuit refinement during development (Carroll et al., 2023; Schafer et al., 2012; Stevens et al., 2007), and has been implicated in several neurodegenerative diseases (Bourel et al., 2021; Hernandez-Encinas et al., 2016; Litvinchuk et al., 2018). In the auditory system, complement signaling has been shown to support auditory function under normal physiological conditions (Brown et al., 2023; Qi et al., 2021). However, the functional role of complement proteins in the cochlea during pathological conditions such as NIHL remains unclear.

The complement system consists of numerous complement proteins and regulatory molecules that interact through three major pathways—the classical, lectin, and alternative pathway (Dunkelberger & Song, 2010; Fishelson et al., 2001; Ricklin et al., 2016; Sarma & Ward, 2011; Vandendriessche et al., 2021). Each pathway can be triggered by distinct pathogen- or damage-associated molecular patterns, yet all converge on the formation of C3 convertases—enzyme complexes that activate and cleave C3 into its active fragments, C3a and C3b (Dunkelberger & Song, 2010; Fishelson et al., 2001; Ricklin et al., 2016; Sarma & Ward, 2011; Vandendriessche et al., 2021). C3a serves as a potent chemoattractant that recruits immune cells and modulates inflammation, whereas C3b functions as an opsonin that tags pathogens and damaged host cells for clearance (Dunkelberger & Song, 2010; Fishelson et al., 2001; Ricklin et al., 2016; Sarma & Ward, 2011; Vandendriessche et al., 2021). While the complement system involves a wide array of proteins, C3 is a particularly compelling target for investigation as a convergence point for all pathways and as a functional mediator of immune responses.

In this study, we investigated the function of C3 in the cochlea during NIHL and found that C3 is activated after noise injury, selectively targeting regions of OHC loss, orphan MOC efferent boutons, and residual cellular debris. Its activation is spatiotemporally dynamic across different structures, highlighting the progression of noise-induced tissue damage and clearance at the auditory sensory epithelium. We further demonstrated that C3 is also recruited during IHC-specific degeneration and cochlear synaptopathy. Finally, we showed that while C3 is dispensable for normal hearing and the development of NIHL, including the extent of OHC degeneration at 14 days after severe noise trauma, lack of C3 prevented MOC efferent bouton degeneration. Together, our findings reveal a damage-selective, temporally dynamic activation of C3 in noise-induced cochlear pathology, which strongly suggests that complement activation plays important roles in debris clearance and synaptic remodeling in the injured cochlea.

## Materials and Methods

### Animals

All experiments were conducted under the guidelines of the protocols approved by the Institutional Animal Care and Use Committee of The Ohio State University (IACUC protocol number: 2018A00000055-R2). Mice used in this study were maintained under a 12:12-h light/dark cycle and had ad libitum access to food and water. Ambient noise level was measured inside the facility and was less than 70 dB SPL in broadband, and below 30 dB SPL at 10 kHz. CBA/CaJ mice, C57BL/6 mice, and homozygous C3-deficient mice (C3−/−; line B6.129S4-C^3tm1Crr^/J; JAX: 003641) of either sex, were purchased from The Jackson Laboratory (Bar Harbor, ME, USA), bred, and maintained at the animal facility at The Ohio State University (Columbus, OH, USA).

### Generation of Calb2-iDTR Mice

Homozygous Calb2-Cre mice (B6(Cg)-Calb2*^tm1(cre)Zjh^*/J; JAX: 010774) and homozygous ROSA26iDTR knock-in mice (C57BL/6-*Gt(ROSA)26Sor^tm1(HBEGF)Awai^*/J; JAX: 007900) were purchased from The Jackson Laboratory (Bar Harbor, ME, USA), and were crossed to create Calb2-iDTR mice that express diphtheria toxin (DT) receptors in calb2-expressing IHCs, which are susceptible to ablation following DT injection. Mice at 11-17 weeks of age received two intraperitoneal DT injections at the dose of 20ng/g on two consecutive days, and their cochleae were harvested one day after the second injection.

### Auditory Brainstem Response (ABR)

Auditory brainstem responses (ABRs) to clicks and tone bursts of various frequencies were recorded to assess hearing status in mice, following protocols previously described (Wang et al., 2019). Briefly, mice were deeply anesthetized using ketamine (100 mg/kg) and xylazine (10 mg/kg) via IP injection, and positioned within a sound-attenuating chamber. Core body temperature was maintained at approximately 36 °C using a feedback-controlled heating pad.

ABRs to sound stimuli were recorded using the RZ6-A-P1 system and BioSigRZ software (Tucker-Davis Technologies). Monophasic clicks (0.1 ms duration, 21 presentations per second, alternating phase) were delivered via a free-field MF1 magnetic speaker placed 10 cm from the ipsilateral ear. Needle electrodes were inserted at the ipsilateral pinna and vertex, with the ground electrode placed at the rump. For each sound level, responses were averaged over 512 repetitions.

### Noise Exposure

Mice at the age of 8-10 weeks were placed in a wire mesh cage suspended in a reverberant exposure box, unanesthetized and unrestrained, and exposed to an octave band (8 – 16 kHz) noise for two hours at the level of 112 dB SPL. Mice were euthanized at different time points ranging from one hour to 60 days post exposure to examine cochlea damage across time.

### Tissue Isolation and Whole Mount Cochlea Preparation

Under deep anesthesia following ABR measurement, mice were decapitated, and the skulls were opened to retrieve the temporal bones for subsequent processing. Each cochlea was carefully dissected out of the temporal bone and flushed with 4% paraformaldehyde (PFA) in 0.1 M phosphate buffered saline (PBS). Cochleae were fixed overnight in ice-cold 4% PFA in 0.1 M PBS and then decalcified using 0.12 M EDTA in 0.1 M PBS for 48 hours. Apex, middle and base turns of the cochlea were dissected out of the labyrinth under a dissection microscope in 0.1 M PBS, followed by immunohistochemistry as detailed below.

### Immunohistochemistry

After tissue preparation, whole-mount cochlea tissue was stained as free-floating sections in a 12-well plate. Tissue sections were blocked using 2% BSA, 0.1% Triton X-100 in 0.1 M PBS for 1 h at room temperature. Following blocking, primary antibodies were applied overnight at 4°C in 1% BSA, 0.1% Triton X-100 in 0.1 M PBS. Primary antibodies used were: guinea pig anti-VAChT (#139105, Synaptic Systems, dilution 1:500), guinea Pig anti-Homer1 (#160005, Synaptic Systems, dilution 1:500), mouse anti-CtBP2 (#612044, BD Biosciences, dilution 1:500), rat anti-C3 complement (#ab11862, Abcam, dilution 1:500), rabbit anti-Myosin-VIIA (#25-6790, Proteus Biosciences, dilution 1:500). Sections were then rinsed three times with 0.1 M PBS for 15 min at room temperature, then secondary antibodies: donkey anti-mouse IgG, Alexa Fluor 488 (#A21202, Thermo Fisher Scientific, dilution 1:250), donkey anti-rat IgG, Alexa Fluor 488 (#A21208, Thermo Fisher Scientific, dilution 1:250), goat anti-rat IgG, Alexa Fluor 594 (#A11007, Thermo Fisher Scientific, dilution 1:250), goat anti-rabbit IgG, Alexa Fluor 594 (#A11037, Thermo Fisher Scientific, dilution 1:250), goat anti-rabbit IgG, Alexa Fluor 750 (#A21039, Thermo Fisher Scientific, dilution 1:250), goat anti-guinea pig IgG, Alexa Fluor 647 (#A21450, Thermo Fisher Scientific, dilution 1:250), were applied overnight at 4°C in 1% BSA, 0.1% Triton X-100 in 0.1 M PBS. Finally, DAPI Fluoromount-G mounting medium (Southern Biotech) was used to mount the slices and highlight the cell nucleus.

### Imaging

All tissues were imaged using an FV3000 confocal microscope (Olympus, Tokyo, Japan). Wide-view images were collected using a 10×air objective, a z-step of 2 μm, and a resolution of 1,024 ×1,024. High-magnification images were collected using a 60×oil immersion objective with 2.0 digital zoom, a z-step of 0.7 or 0.3 μm and resolution of 800 × 800.

### Image Analysis

Image processing and analysis was conducted using ImageJ (U. S. National Institutes of Health, Bethesda, MD, USA) software and Imaris software (version 9.5.0; Oxford Instruments). Using ImageJ software, Z-stack volumetric confocal images were reduced to 2D maximum projections and individual channels were quantified as appropriate for stained area (VAChT-labeled MOC efferent boutons) and mean fluorescent intensity (C3; VAChT-labeled MOC efferent bouton) within regions of interest (ROI) following supervised auto-thresholding. Colocalization of C3 proteins with CtBP2-labeled ribbon puncta and/or Homer1-labeled auditory nerve boutons were analyzed with ImageJ.

Three-dimensional reconstruction of confocal images was performed using Imaris software. Three-dimensional reconstruction of C3 protein (green), Myosin-VIIA-labeled sensory hair cells (red), and VAChT-labeled MOC efferent boutons (magenta) that synapse onto the base of outer hair cells were made using semi-automated Imaris tools, including masking uninterested areas and rendering surfaces in three separate channels. The basal region of the Myosin-VIIA-labeled outer hair cell that contains nucleus and contacts MOC efferent boutons were not included when rendering surfaces, to prevent overlapping across channels for better visualization. Opsonization by C3 protein was determined based on the colocalization pattern of C3 protein with other cellular markers.

### Statistics

All data are reported as the mean ± SD, unless otherwise noted. Statistical analyses were performed using GraphPad Prism 10 software. One-way ANOVA and following Tukey’s multiple comparisons test were used to compare means of more than two groups. Two-way ANOVA was used to make comparisons across more than two groups followed by Šídák’s or Tukey’s multiple comparisons test. All statistic tests were two-tailed unpaired, and *P* -values are denoted as: ns (not significant) = *P* > 0.05, \**P* < 0.05, \*\**P* < 0.01, \*\*\**P* < 0.001, and \*\*\*\**P* < 0.0001.

## Results

### Noise injury induces an upregulation of C3 activity that selectively targets damaged regions of the organ of Corti in mice with NIHL

Tissue damage can activate complement protein C3, which opsonizes and promotes the phagocytosis of endogenous debris and triggers inflammatory signaling through the release of proinflammatory mediators (Fishelson et al., 2001; Ricklin et al., 2016; Vandendriessche et al., 2021). It is widely studied that noise overexposure causes tissue injury and cell death in the cochlea underlying NIHL (Wang et al., 2002). However, the production, activation, and the roles of C3 in the cochlea after noise injury remain unclear. To address this knowledge gap, we examined the involvement of C3 in NIHL.

Young (8-10 weeks of age) CBA/CaJ mice were exposed to an octave band noise (8–16 kHz) at 112 dB SPL for 2 hours. Such high-level NE was reported to induce a permanent threshold shift in CBA/CaJ mice (Wang et al., 2002). Consistently, we observed severe hearing loss as shown by significant elevation of auditory brainstem response (ABR) threshold to clicks and tone bursts at 8, 12, 16, and 32 kHz, as well as significantly reduced ABR wave I amplitude to clicks in noise-exposed (NE) animals 14 days post-NE (Fig. 1A, B). Histological analysis revealed region-specific cochlear pathology. Compared with the control group, NE mice showed a significant loss of outer hair cells (OHCs), and a marked reduction in ribbon counts between inner hair cells (IHCs) and spiral ganglion neurons (SGNs), but only in the basal turn of the cochlea (Fig. 1C-E, 1N-P). In contrast, IHCs themselves remained largely intact, with no significant reduction in number (Fig. 1D-E, Q). No significant structural changes were observed in the apical or middle cochlear turns (Fig. 1C-E, 1N-Q). Together, these findings showed that our noise-exposure protocol induced significant ABR threshold shifts, region specific sensory hair cell death, and cochlear synaptopathy.

**Fig 1.**
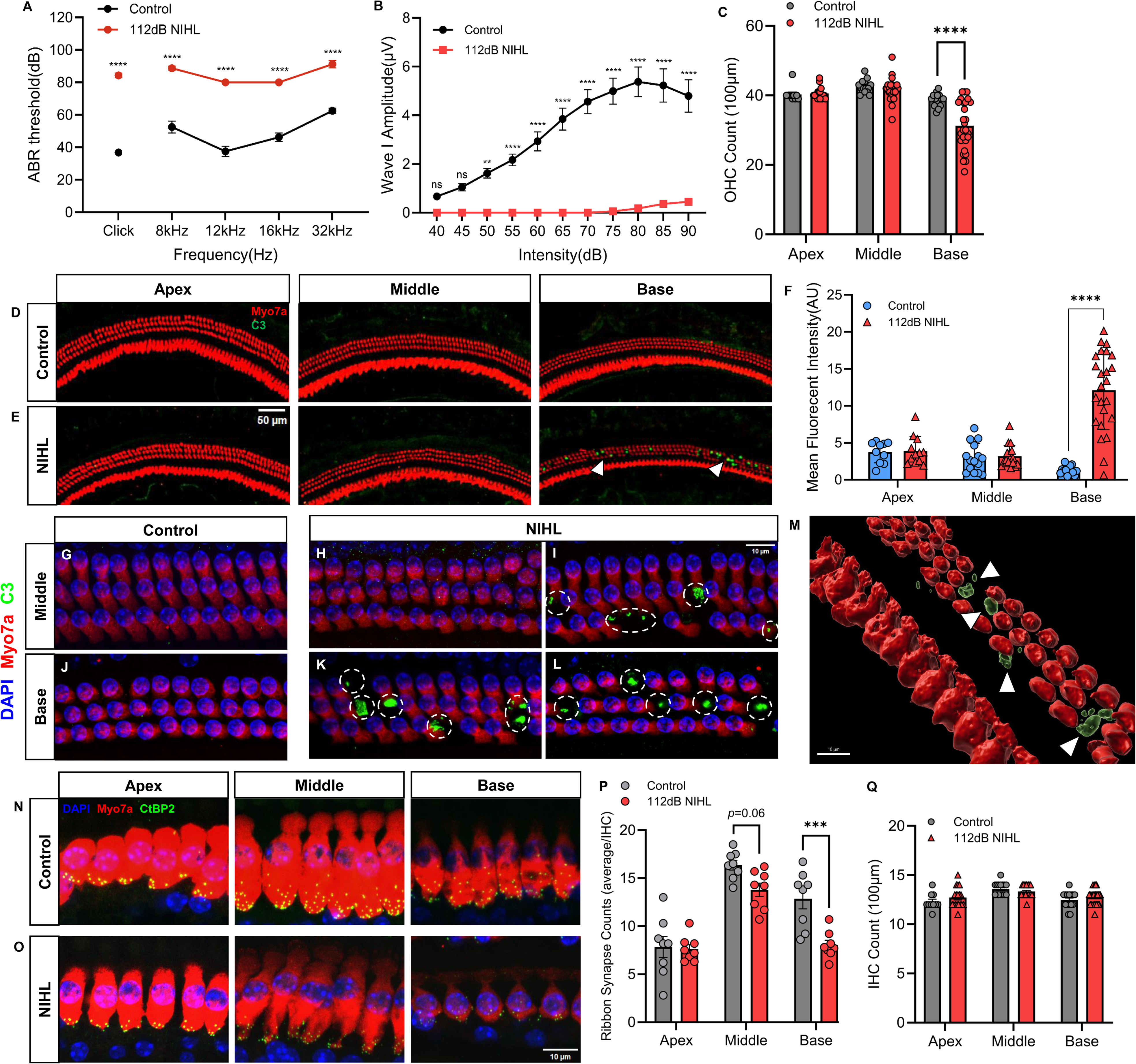
C3 selectively targets damaged regions of the organ of Corti in mice with noise-induced hearing loss (NIHL). **A** ABR thresholds in response to clicks and tone bursts at 8, 12, 16, and 32 kHz in control mice (n=5) and noise-exposed mice (n=8). ****, *P* < 0.0001, two-way ANOVA followed by Šídák’s multiple comparisons test. **B** Growth curves of ABR click wave I amplitude in both control (n=5) and exposed mice (n=8). ****, *P* < 0.0001, two-way ANOVA followed by Šídák’s multiple comparisons test. **C** The number of OHCs from apical, middle, and basal turns of control and noise-exposed cochleae (n≥10 regions of interest from independent confocal images for each group). ****, *P* < 0.0001, two-way ANOVA followed by Šídák’s multiple comparisons test. **D-E** Representative immunofluorescent staining images showing the DAPI (blue) labeled nuclei, myosin 7a (red) labeled sensory hair cells, and C3 proteins (green) in apical, middle, and basal turns of control and noise-exposed cochleae. C3 was found to predominantly accumulate in the damaged OHC regions in the basal turns (white arrowheads). **F** Mean C3 fluorescent intensity of OHC layers in apical, middle, and basal turns of control and noise-exposed cochleae (n≥10 regions of interest from independent confocal images for each group). ****, *P* < 0.0001, two-way ANOVA followed by Šídák’s multiple comparisons test. **G-L** High magnification immunofluorescent staining images of DAPI (blue) labeled nuclei, myosin 7a (red) labeled OHCs, and C3 proteins (green) in middle and basal turns of control and noise-exposed cochleae. (**H**) No C3 activity was found in intact middle turns. C3 was found to predominantly accumulate in the gaps left by missing outer hairs in severely damaged basal turns (**K-L**) and occasionally impaired middled turns (**I**) (dashed circles). **M** 3D reconstruction of representative immunofluorescent staining image (**K**), showing myosin 7a-labeled sensory hair cells (red) and C3 proteins (green). **N-O** Representative immunofluorescent staining images showing the CtBP2 (green) labeled ribbons and myosin 7a (red) labeled inner hair cells in apical, middle, and basal turns of control and noise-exposed cochleae. **P** The number of inner hair cells from apical, middle, and basal turns of control and noise-exposed cochleae (n>10 regions of interest from independent confocal images for each group). ****, *P* < 0.0001, two-way ANOVA followed by Šídák’s multiple comparisons test. **Q** The average number of ribbons counts per inner hair cell from apical, middle, and basal turns of control and noise-exposed cochleae (n=8 per group). ***, *P* < 0.001, two-way ANOVA followed by Šídák’s multiple comparisons test. (**A-C, F, P-Q**) All Data were shown as mean ±SEM. (**A-Q**) All noise-exposed mice were examined 14 days post NE.

To determine whether C3 is activated in response to noise-induced cochlear damage, we examined C3 activity using immunofluorescent staining. Under baseline condition, C3 antibody labeling was minimal across all cochlear turns, indicating low constitutive activity (Fig. 1D). In contrast, 14 days after NE, we observed robust C3 accumulation selectively in regions of OHC loss within the basal turns (Fig. 1E). Notably, no C3 labeling was detected in IHC regions, which remained structurally intact after NE (Fig. 1E), or in the apical and middle turns of the cochlea, where no OHC loss was observed (Fig. 1C). In NE animals, we also detected C3 staining within blood vessels in the stria vascularis and spiral limbus, suggesting a systemic origin of C3 protein. To quantify C3 activation, we measured mean C3 fluorescence intensity in the organ of Corti across cochlear turns. C3 levels remained low in the well-preserved apical and middle turns but were significantly elevated in the severely noise-damaged basal turns of NE animals (Fig. 1F), suggesting a damage-dependent pattern of activation.

To further examine the spatial localization of C3, we acquired high-magnification confocal images of OHC regions at middle and basal turns in control and NE cochleae (Fig. 1G-L). In NE animals, large C3 deposits were located specifically in the gaps left by missing OHCs (Fig. 1I, K, L). Our NE protocol is known to spare the apical and middle turns from significant pathology (Fig. 1C, P, Q), which aligns with the absence of C3 activation in these regions (Fig. 1G, H).

However, in cases where occasional OHC loss did occur in the middle turn, similar C3 accumulation was observed (Fig. 1I), reinforcing the damage-dependent nature of C3 activity. Three-dimensional reconstructions of representative images confirmed that C3 deposits precisely occupied the spaces formerly occupied by OHCs (Fig. 1M). These data collectively demonstrate that C3 activity was tightly linked to tissue damage and selectively targeted degenerating OHC regions in the organ of Corti at 14 days post-NE.

### C3 opsonizes impaired orphan MOC efferent boutons with heterogeneous morphology

One of the major functions of C3 is to opsonize impaired cells and synapses, facilitating their recognition and clearance by phagocytic cells (Fishelson et al., 2001; Ricklin et al., 2016; Vandendriessche et al., 2021). Having established that C3 is involved in noise-induced pathology in a damage-dependent manner, we next sought to identify the specific cellular targets of C3 within regions where OHCs had already degenerated. Since these cells were no longer present at 14 days post-NE, we focused on persistent cellular elements that may remain in their absence.

Previous studies suggested that medial olivocochlear (MOC) efferent boutons, which normally synapse onto OHCs, can persist at the sites of missing OHCs for an extended period following OHC death in age-related hearing loss (Dörje et al., 2024; Fu et al., 2010). We hypothesized that a similar process may occur following NE, with C3 potentially opsonizing these "orphaned" and dysfunctional MOC boutons that need to be cleared. Consistent with our hypothesis, we stained MOC terminals using VAChT antibody and observed orphan MOC efferent boutons in the gaps left by degenerated OHCs in the basal turns, 14 days after NE (Fig. 2A-E). These boutons colocalized with C3 deposits, suggesting that they are a major target of C3 opsonization at this time point. 3D reconstructions of immunofluorescent images revealed that C3 encapsulated these boutons (Fig. 2H-M), suggesting active opsonization. A small subset of orphan boutons lacked C3 labeling (Fig. 2A, B, N), likely representing MOC boutons not yet affected by ongoing tissue degeneration, reflecting a progressive pattern of pathology in the organ of Corti.

**Fig 2.**
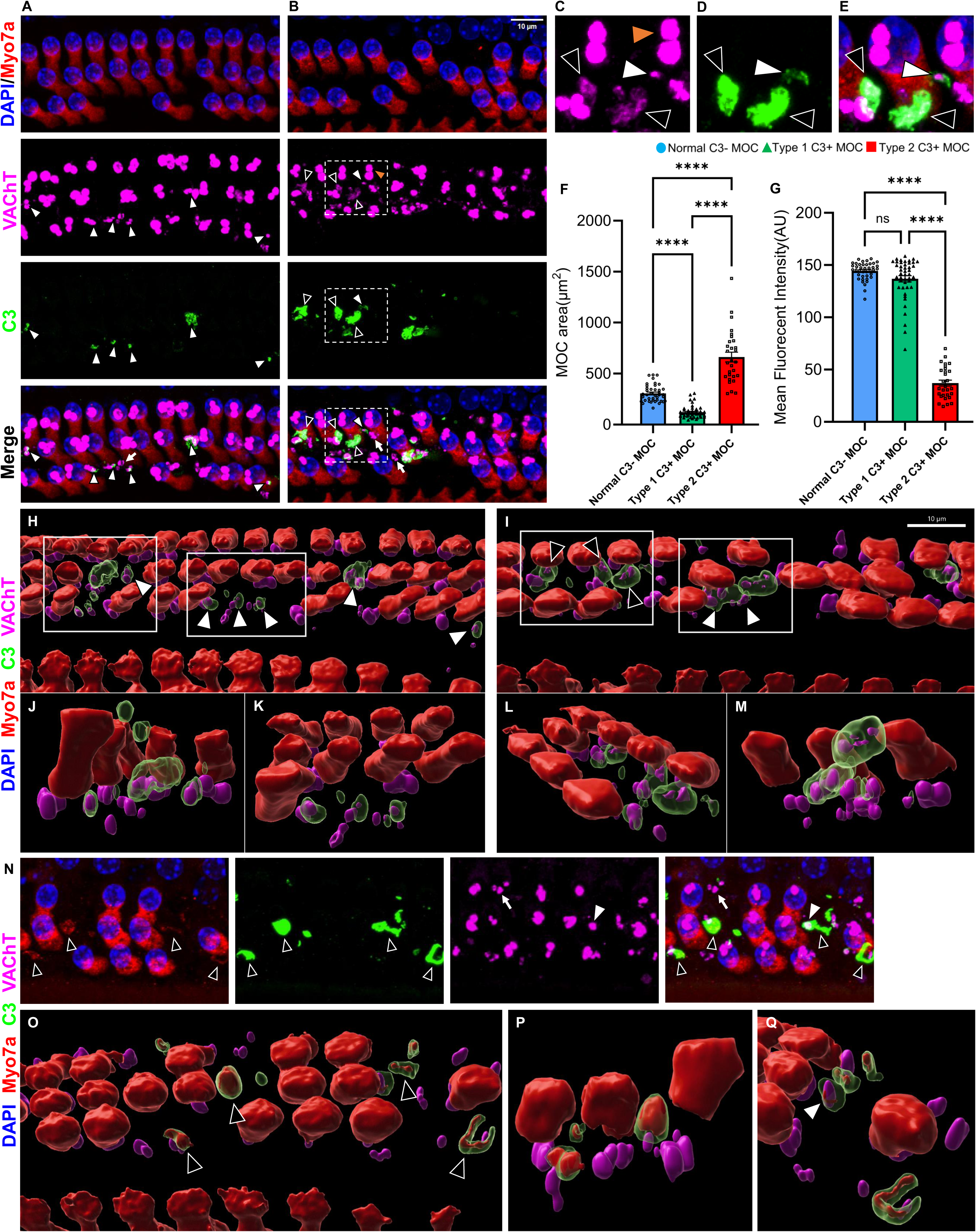
C3 opsonizes impaired orphan MOC nerve terminal boutons of missing OHCs. **A-B** Representative immunofluorescent staining images showing the DAPI (blue) labeled nuclei, myosin 7a (red) labeled sensory hair cells, VaChT (magenta) labeled MOC nerve terminal boutons, and C3 proteins (green) of noise-exposed cochleae. C3 strongly colocalizes with orphan MOC nerve terminal boutons (white and black arrowheads) located in the gaps of missing OHCs. **C-E** Magnified views of regions of interest indicated in (**B**). C3 positive orphan MOC nerve terminal boutons showed heterogeneous morphology: type1 (white arrowheads) and type2 (black arrowheads), that differ from normal C3 negative MOC nerve terminal boutons (orange arrowhead). **F** Comparisons of mean areas of normal, type 1 and type 2 MOC nerve terminal boutons (n≥29 MOC nerve terminal boutons for each group). ****, *P* < 0.0001, one-way ANOVA followed by Tukey’s multiple comparisons test. **G** Comparisons of mean VaChT fluorescent intensities of normal, type 1 and type 2 MOC nerve terminal boutons (n≥29 MOC nerve terminal boutons for each group). ns, nonsignificant; ****, *P* < 0.0001, one-way ANOVA followed by Tukey’s multiple comparisons test. **H-I** 3D reconstructions of representative immunofluorescent staining images (**A&B**) of myosin 7a (red) labeled sensory hair cells, VaChT (magenta) labeled MOC nerve terminal boutons, and C3 proteins (green). C3 opsonizes impaired orphan MOC nerve terminal boutons (white and black arrowheads) located in the gaps of missing OHCs. **J-K** Magnified views of region of interests indicated in (**H**). **L-M** Magnified views of region of interests indicated in (**I**). **N** Representative immunofluorescent staining images showing the DAPI (blue) labeled nuclei, myosin 7a (red) labeled sensory hair cells, VaChT (magenta) labeled MOC nerve terminal boutons, and C3 proteins (green) of noise-exposed cochleae. C3 colocalizes with MOC nerve terminal boutons (white arrowhead), and with myosin 7a positive OHC debris (black arrowheads). **O** 3D reconstruction of representative immunofluorescent staining images of myosin 7a (red) labeled sensory hair cells, VaChT (magenta) labeled MOC nerve terminal boutons, and C3 proteins (green). C3 opsonizes orphaned MOC nerve terminal boutons (white arrowhead) and myosin 7a positive OHC debris (black arrowheads), located in the gaps of missing OHCs. **P-Q** Magnified views of C3-opsonized orphan MOC nerve terminal boutons (white arrowhead) and myosin 7a positive OHC debris (black arrowheads) indicated in (**N, O**). (**A-B, N**) C3-negative orphan MOC nerve terminal boutons of missing OHCs were also observed (white arrows). (**F-G**) All Data were shown as mean ±SEM. (**A-Q**) All noise-exposed mice were examined 14 days post NE.

We then noted that putative C3-opsonized boutons exhibit heterogeneous morphologies that differed markedly from healthy MOC boutons (Fig. 2C-E). Based on morphology, we categorized these C3-opsonized MOC boutons into two types: type 1 boutons appeared shrunken, while type 2 were dispersed and swollen (Fig. 2C-E). These morphological patterns resemble those reported in prior NE studies (Boero et al., 2018; Canlon et al., 1999). Quantitative analysis showed that type 1 boutons had significantly smaller areas but retained normal fluorescence intensity (Fig. 2F-G), suggesting structural atrophy without loss of marker expression. In contrast, type 2 boutons were significantly larger than both type 1 and normal C3-negative boutons and exhibited reduced fluorescence intensity (Fig. 2F-G), indicating disintegration. The exact degeneration process of MOC boutons after NE remains unclear. These distinct morphologies may reflect different stages or mechanisms of MOC degeneration, possibly linked to varied OHC death pathways.

In addition to the opinionized MOC boutons, we also occasionally observed Myo7a-positive fragments at the sites of OHC loss, which co-localized with C3 (Fig. 2N). These fragments resemble prestin-positive OHC remnants described in previous studies (Abrashkin et al., 2006; Yu et al., 2011) and likely represent the debris of incompletely degraded OHCs. 3D reconstructions of representative immunofluorescent staining images confirmed that C3 not only targeted orphan MOC boutons but also encapsulated these residual OHC fragments (Fig. 2O-Q). Together, these findings suggest that C3 plays a key role in targeting degenerating cellular remnants in the noise-damaged cochlea. Specifically at 14 days post-NE, a relatively chronic time point when OHC degeneration is largely complete, C3 is activated to opsonize remaining orphaned MOC boutons and remaining OHC debris.

### Dynamics of C3 opsonization in the organ of Corti during noise-induced pathology

Noise-induced cochlear damage unfolds progressively, affecting distinct traumatized cellular structures over time (Hu et al., 2002; Wang et al., 2002). To better understand how C3 activation evolves during this process, we analyzed its spatial and temporal dynamics at multiple time points following NE. Additional young CBA/CaJ mice (8–10 weeks old) were exposed to the same noise protocol (8–16 kHz at 112 dB SPL for 2 hours), and cochlear tissues were collected at 1 hour (D0), 1 day (D1), 5 days (D5), and 60 days (D60) post-exposure. ABR thresholds showed significant elevations to both click and tone burst stimuli at 8, 12, 16, and 32 kHz at D0, D5, and above reported 14 days (D14) post NE (Fig. 3A). Partial recovery from acute deafness was observed between D0 and D5 at low and middle frequency hearing, but no further improvements were seen beyond D5, indicating the establishment of permanent hearing loss (Fig. 3A).

**Fig 3.**
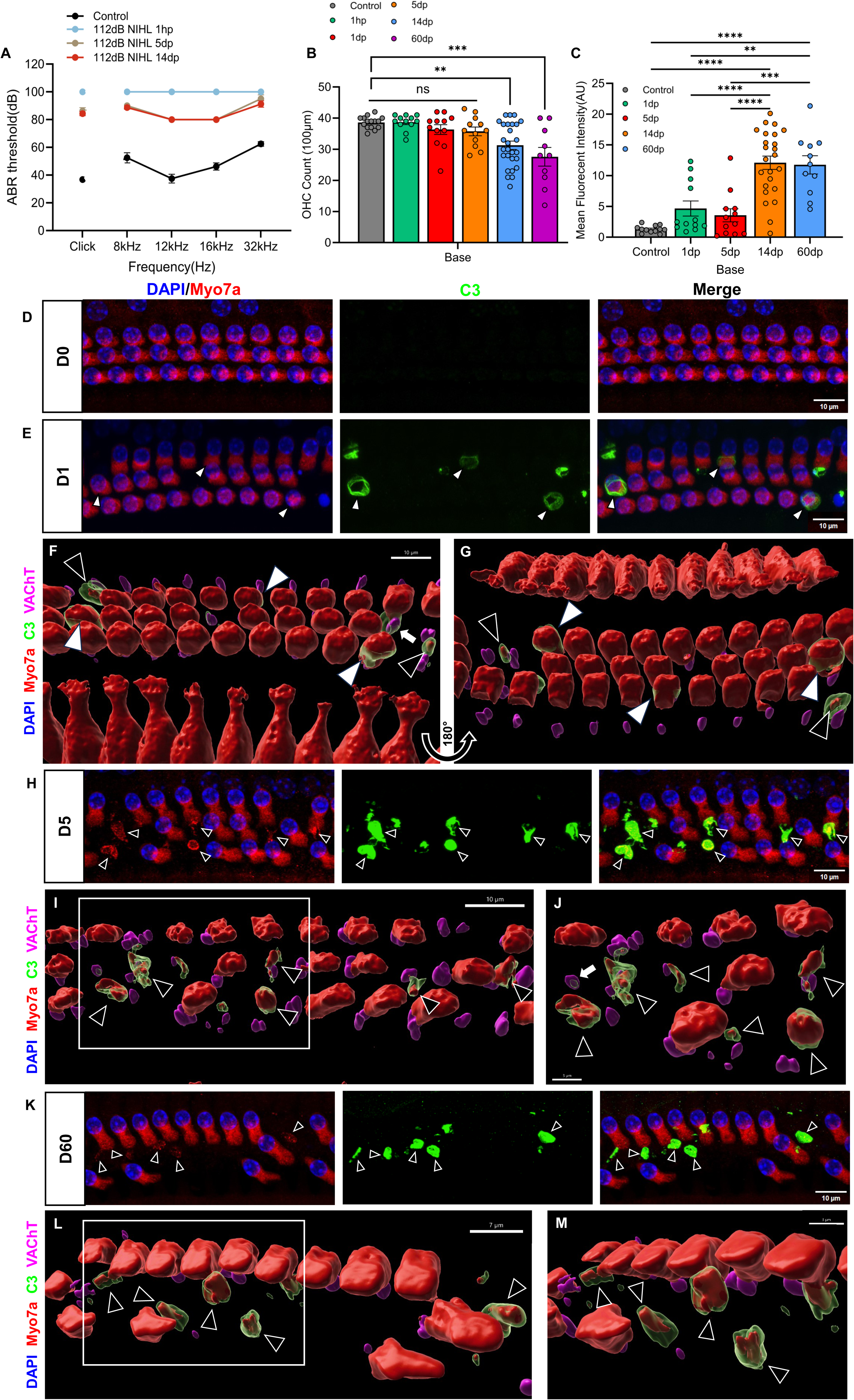
C3 opsonizes various degenerating cellular structures during the progression of noise-induced pathology. **A** ABR thresholds in response to clicks and tone bursts at 8, 12, 16, and 32 kHz in control mice (n=5) and noise-exposed mice at 1 hour (1h; n=4), 1 day (D1; n=4), 5 days (D5; n=4), and 14 days (D14; n=8) post NE. Significant effects were analyzed with one-way ANOVA followed by Šídák’s multiple comparisons test. **B** The number of OHCs in basal turns of control and noise-exposed cochleae from different post-exposure time points (n≥10 regions of interest from independent confocal images for each group). ns, nonsignificant; ***, *P* < 0.001; **, *P* < 0.01, one-way ANOVA followed by Šídák’s multiple comparisons test. **C** Mean C3 fluorescent intensity of OHC layers in basal turns of control and noise-exposed cochleae from different post-exposure time points (n≥11 regions of interest from independent confocal images for each group). ****, *P* < 0.0001; ***, *P* < 0.001; **, *P* < 0.01, one-way ANOVA followed by Šídák’s multiple comparisons test. **D** Representative immunofluorescent staining images showing the DAPI (blue) labeled nuclei, myosin 7a (red) labeled sensory hair cells, and C3 proteins (green) in the basal turn 1h post NE. Minimal C3 activity was observed 1h post NE. **E** Representative immunofluorescent staining images showing the DAPI (blue) labeled nuclei, myosin 7a (red) labeled sensory hair cells, and C3 proteins (green) in the basal turn 1 day post NE. C3 opsonizes damaged OHCs that potentially just start degeneration (white arrowheads). **F-G** 3D reconstructions of representative immunofluorescent staining image (**E**), showing myosin 7a (red) labeled sensory hair cells, VaChT (magenta) labeled MOC nerve terminal boutons, and C3 proteins (green). Additional to damaged OHCs (white arrowheads), C3 also opsonizes a few smaller myo7a positive debris (black arrowheads) of degenerating OHCs. **H** Representative immunofluorescent staining images showing the DAPI (blue) labeled nuclei, myosin 7a (red) labeled sensory hair cells, and C3 proteins (green) in the basal turn 5 days post NE. C3 predominantly opsonizes OHC debris as damaged OHCs continued to degenerate (black arrowheads). **I** 3D reconstructions of representative immunofluorescent staining image (**E**), showing myosin 7a (red) labeled sensory hair cells, VaChT (magenta) labeled MOC nerve terminal boutons, and C3 proteins (green). **J** Magnified view of region of interest indicated in (**I**). **K** Representative immunofluorescent staining images showing the DAPI (blue) labeled nuclei, myosin 7a (red) labeled sensory hair cells, and C3 proteins (green) in the basal turn 60 days post NE. C3 predominantly opsonizes OHC debris as damaged OHCs continued to degenerate (black arrowheads). **L** 3D reconstructions of representative immunofluorescent staining image (**K**), showing myosin 7a (red) labeled sensory hair cells, VaChT (magenta) labeled MOC nerve terminal boutons, and C3 proteins (green). **M** Magnified view of region of interest indicated in (**L**). (**A-C**) All Data were shown as mean ±SEM. (**F, G**) A few C3-opsonized orphan MOC boutons were also observed (white arrows).

Previous studies suggested that OHCs degenerate through apoptosis or necrosis within 24 hours, with degeneration expanding to additional cochlear frequency regions and tapering off by day 5 post-exposure (Hu et al., 2002; Ping Yang et al., 2004). We found OHC counts in the basal turn showed significant OHC loss over time after NE compared to control animals, but in contrast to previous studies, post hoc tests showed that OHC loss was more profound at 14- and 60-days post NE, (Fig. 3B). This discrepancy is likely due to the variation in NE levels and frequency ranges used in earlier studies (Hu et al., 2002; Ping Yang et al., 2004) and here. Significant loss of OHCs was observed at both D14 and D60, with no significant difference between these two time points (Fig. 3B). These results suggest that while many OHCs may have been functionally compromised shortly after exposure, actual cell loss appeared to peak between D5 and D14 and remained stable through D60. We next assessed the change in C3 activity across different post exposure time points, by measuring mean C3 fluorescence intensity within OHC regions. Due to the damage-dependent activity of C3, we focused our analysis on the basal turns of the cochlea, where damage was most severe. Consistent with the progression of OHC loss after NE (Fig. 3B), the overall C3 signal significantly increased over time (Fig. 3C), despite that the increase in mean fluorescent intensity was not statistically significant at D0, D1 to D5 compared to the controls, matching the low levels of OHC death at these acute time points (Fig. 3B). At one-hour post NE, we did not observe any selective C3 labeling in the OHC region (Fig. 3D), where OHC loss was rarely seen. In contrast, we found that at all other time points including D1, D5 and D60 post NE, cochlea regions with OHC loss showed localized C3 deposits (Fig. 3E, H, K). These results further suggest that C3 is activated in the cochlea upon noise injury and dynamically opsonizing damaged cellular structures in the OHC region.

High-magnification and 3D reconstructed representative confocal images further elucidated this timeline. At D0, C3 staining in the organ of Corti was minimal (Fig. 3D), indicating a time delay between immediate tissue damage and C3 activation. By D1, C3 colocalized with damaged but largely intact OHCs that appeared to be undergoing early degeneration (Fig. 3E-G). Small Myo7a-positive fragments—likely debris of ongoing OHC degeneration—were also observed in 3D reconstruction (Fig. 3F-G), although not obvious in the maximum projection of Z-stacked confocal images. At D5, C3 labeling was associated with more OHC debris (Fig. 3H-J). The Myo7a-positive debris observed at D1 and D5 possibly represent a later stage of C3-opsonized OHCs as cells continued to degenerate. Occasional C3-labeled MOC boutons were present at D1 and D5 (Fig. 3F, I, J), but they were rare and not the predominant C3 targets at these early stages. At D60, a time point well beyond the expected resolution of inflammation (Kalinec et al., 2017; Wood & Zuo, 2017), we examined whether C3 activation was eventually resolved and surprisingly found that C3 activity remained elevated (Fig. 3C). Persistent C3 labeling was observed in association with residual OHC debris at D60, like those seen at D14 and D5 (Fig. 3K-M). This sustained complement activation and persistent OHC debris is puzzling given the lack of further OHC loss between D14 and D60 (Fig. 3B). The organ of Corti lacks resident immune cells under normal condition (Hirose et al., 2005; Okano et al., 2008; Yang et al., 2015). Following injury, supporting cells—particularly Deiters’ cells (DCs)—act as ‘amateur’ phagocytes that engulf and clear cellular debris (Abrashkin et al., 2006; Anttonen et al., 2014; Hirose et al., 2017; Monzack et al., 2015). However, their phagocytic capacity could be limited, resulting in incomplete clearance of C3-opsonized material within 60 days post injury and the consequent chronic C3 activation toward those remaining cellular debris. Taken together, these findings demonstrate that C3 activation in the cochlea is both damage-dependent and temporally dynamic. C3 opsonizes distinct targets at different stages of pathology, including damaged OHCs (D1 to D5), OHC debris during degeneration and clearance (D5 to D60 and beyond), and ultimately orphaned MOC boutons (D14 up to D60). The persistence of C3 activity through 60 days post-exposure suggests incomplete immune resolution that may underlie chronic inflammation (Seicol et al, 2022), possibly due to the unique immune-privileged environment of the organ of Corti.

### C3 is involved in diphtheria toxin (DT)-induced inner hair cell death and ribbon synapse degeneration

While our NE treatment led to robust C3 activation in regions of OHC loss, it did not induce significant IHC death or C3 accumulation in IHC regions (Fig. 1N-Q). A significant loss of cochlear synapses was observed after NE (Fig. 1P), but it remains unclear whether C3 activity is involved because cochlear synapse loss is transient upon NE and might not be captured at our selective time points. To test whether C3 activity also contributed to tissue damage in the IHC region, we generated a new mouse model of IHC damage by crossing Calb2-Cre mice with ROSA26-iDTR knock-in mice to create Calb2-iDTR mice, which expresses the diphtheria toxin receptor (DTR) in calretinin-expressing IHCs and type I_a_ SGNs. Immunohistological examination revealed significant IHC loss in DT-injected mice (Fig. 4A-C), confirming effective targeted ablation. To determine whether C3 is recruited during IHC degeneration, we examined its spatial distribution in the organ of Corti. High-magnification images revealed C3 accumulation in regions of IHC loss (Fig. 4D-E). Specifically, C3 colocalized with IHCs undergoing active degeneration (Fig. 4D-E), indicating that C3 activation extends to IHC- targeted injury. Notably, C3 deposits in these regions were smaller and less abundant than those observed in noise-damaged OHC regions at 14 days post-exposure. This difference likely reflects the rapid kinetics and mechanisms of DT-induced cell death, which may have progressed beyond the window of peak C3 activity by the time of tissue collection. Only a few IHCs that were still in the process of degeneration were targeted by C3.

**Fig 4.**
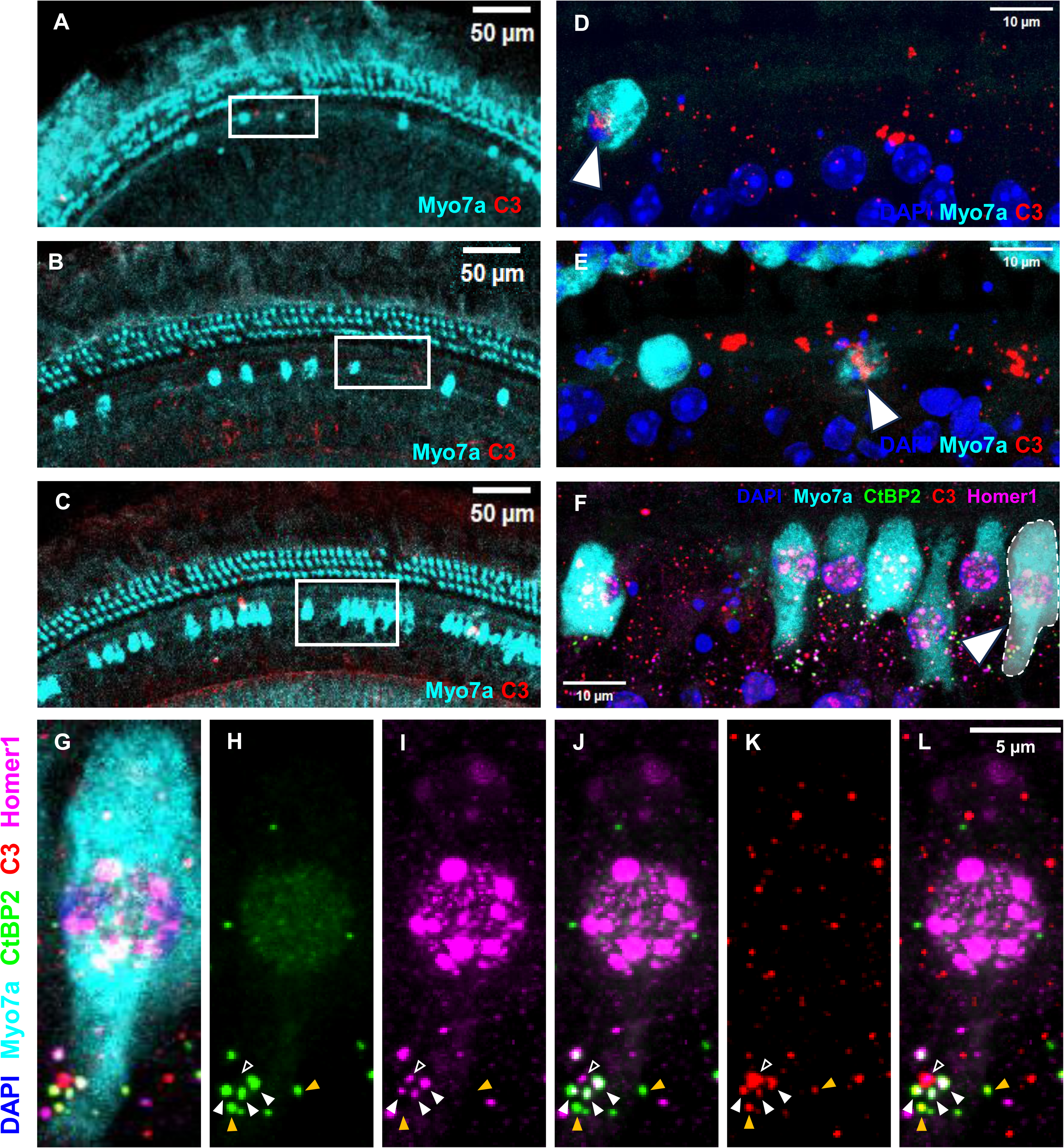
C3 accumulates in damaged IHC regions and opsonizes pre- and/or post synaptic components of degenerating ribbon synapses after Diphtheria toxin (DT) injections. **A-C** Representative immunofluorescent staining images showing the myosin 7a (cyan) labeled sensory hair cells, and C3 proteins (red) in the middle turns of DT injected cochleae. **D-F** High magnification images showing the regions of interest indicated in (**A-C**) respectively. C3 activation was observed in damaged IHC regions (white arrowheads). **G** Representative immunofluorescent staining images showing DAPI (blue) labeled nuclei, myosin 7a (cyan) labeled sensory hair cells, and C3 proteins (red) in the middle turns of DT injected cochleae. Pre- and postsynaptic segments were labeled by CtBP2 (green) and Homer1 (magenta). C3 opsonizes orphan CtBP2 puncta (orange arrowheads), orphan Homer1 puncta (black arrowheads), and CtBP2/Homer1 ribbon synapse pairs (white arrowheads).

We further examined whether C3 also targets synapses between IHCs and SGNs. In addition to IHCs, Calb2-iDTR mice also express DTR in the cochlear synapses of calretinin-expressing type I_a_ SGNs, making both pre- and postsynaptic components of ribbon synapses vulnerable to DT-induced injury. Immunostaining revealed C3 colocalization with orphan presynaptic CtBP2 puncta, orphan postsynaptic Homer1 puncta, and residual CtBP2/Homer1 synapse pairs (Fig. 4F-L), which represent different patterns of synaptic damages from DT treatment. The relatively weaker C3 response observed here was likely due to the abrupt and efficient nature of DT-induced cell death, which limited the temporal window of peak C3 activity. These findings demonstrated that C3 activation is not limited to OHC and MOC-related pathology but also participates in IHC degeneration and cochlear synaptopathy under conditions of targeted injury. It suggests that C3 activation is universally involved in tissue damage in the cochlea, under both NE and DT induced injuries.

### Depleting C3 prevents the degeneration of MOC efferent nerve boutons, but does not significantly impact hearing after severe NIHL

To assess the functional role of C3 activation in noise-induced cochlear pathology, we investigated whether C3 depletion could alter hearing outcomes or cellular degeneration following noise injury using homozygous C3 knockout mice (C3KO) and age-matched wild-type (WT) C57BL/6 mice. Both strains carry the Cdh23*^ahl^* allele that is associated with early-onset hearing loss but show relatively normal hearing at young age (<20 weeks old) (Ison et al., 2007; Mock et al., 2016). In this experiment, young (8–10 weeks of age) WT and C3KO mice were exposed to the same 112 dB SPL octave-band noise (8–16 kHz) for 2 hours to induce NIHL and compared to non-exposed controls. We evaluated the hearing status of these mice by measuring ABRs at 14 days post exposure and collected cochleae to examine tissue damage. At baseline, C3KO mice displayed normal auditory function. ABR thresholds, wave I amplitudes, and OHC counts were comparable to those of WT controls (Fig. 5A-E), indicating that C3 is not required for normal cochlear development or hearing function.

**Fig 5.**
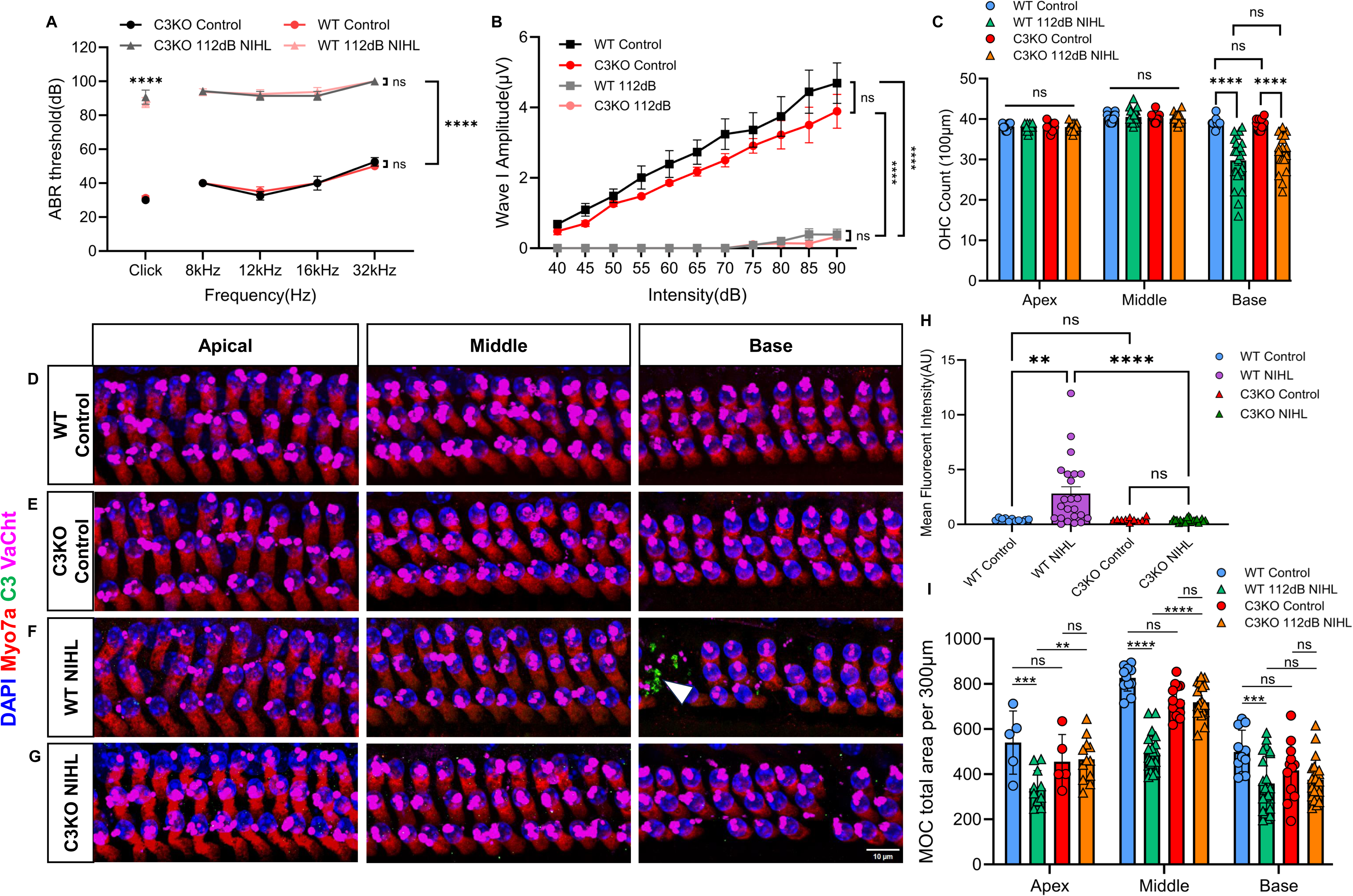
Depleting C3 prevents the degeneration of MOC boutons, but does not affect ABR thresholds and ABR click wave I amplitudes after high intensity noise damage. **A** ABR thresholds in response to clicks and tone bursts at 8, 12, 16, and 32 kHz in control wild-type mice (n=4), control C3-deficient mice (n=4), NE wild-type mice (n=8), and NE C3-deficient mice (n=8). ns, nonsignificant; ****, *P* < 0.0001, two-way ANOVA followed by Šídák’s multiple comparisons test. **B** Growth curves of ABR click wave I amplitude in control wild-type mice (n=4), control C3-deficient mice (n=4), NE wild-type mice (n=8), and NE C3-deficient mice (n=8). ns, nonsignificant; ****, *P* < 0.0001, two-way ANOVA followed by Šídák’s multiple comparisons test. **C** The number of OHCs from apical, middle, and basal turns of cochleae from control wild-type mice, control C3-deficient mice, NE wild-type mice, and NE C3-deficient mice (n≥8 regions of interest from independent confocal images for each group). ns, nonsignificant; ****, *P* < 0.0001, two-way ANOVA followed by Šídák’s multiple comparisons test. **D-G** Representative immunofluorescent staining images showing the DAPI (blue) labeled nuclei, myosin 7a (red) labeled sensory hair cells, and C3 proteins (green) in apical, middle, and basal turns of cochleae from control wild-type mice (**D**), control C3-deficient mice (**E**), NE wild- type mice (**F**), and NE C3-deficient mice (**G**). C3 activation was only found in NE wild-type mice (white arrowhead). **H** Mean C3 fluorescent intensity of OHC layers in the basal turns of cochleae from control wild-type mice, control C3-deficient mice, NE wild-type mice, and NE C3-deficient mice (n≥11 regions of interest from independent confocal images for each group). ns, nonsignificant; ****, *P* < 0.0001; **, *P* < 0.01, one-way ANOVA followed by Tukey’s multiple comparisons test. **I** Total area of MOC nerve terminal boutons per 300 µm from apical, middle, and basal turns of cochleae from control wild-type mice, control C3-deficient mice, NE wild-type mice, and NE C3-deficient mice (n≥5 regions of interest from independent confocal images for each group). ****, *P* < 0.0001; ***, *P* < 0.001; **, *P* < 0.01, two-way ANOVA followed by Šídák’s multiple comparisons test.

14 days following NE, both WT and C3KO mice exhibited severe hearing impairments, as shown by elevated ABR thresholds and reduced wave I amplitudes (Fig. 5A, B). Significant OHC loss was observed in the basal turn of both genotypes (Fig. 5C, F, G). However, no significant differences were found between WT and C3KO mice after NE (Fig. 5A-C). These findings suggest that C3 deletion does not prevent sensory hair cell loss or functional hearing deficits after severe NIHL. To confirm the depletion of C3 activity, we quantified mean C3 fluorescence intensity in the basal turn, where damage was most severe. In WT mice, C3 levels were significantly elevated following NE compared to unexposed WT controls (Fig. 5F, H). In contrast, C3KO mice showed no such increase (Fig. 5G, H), confirming effective loss of C3 function by gene ablation. In the central nerve system, ablating C3 prevents age-related hippocampal neuronal decline (Shi et al., 2015), suggesting that C3 may be detrimental to cellular pathology. However, we found no protective effect in C3KO mice on ABR thresholds or OHC survival, possibly due to the ceiling effect that the overwhelming severity of injury caused by the 112 dB SPL exposure may have masked any protective effects of C3 depletion.

We next examined the effect of C3 depletion on the degeneration of MOC efferent boutons by measuring the reduction in MOC bouton size. Given the variability in individual bouton size, we quantified total bouton area per 300 μm section across cochlear turns, rather than average individual bouton size. At baseline, C3KO mice displayed normal MOC bouton areas across all cochlear regions (Fig. 5D, E, I). After NE, WT mice showed a significant reduction in total MOC bouton area in noise-damaged basal turns, and even in apical and middle regions without OHC loss (Fig. 5C, D, F, I), indicating that noise-induced MOC degeneration can occur independently of OHC death. Strikingly, this degeneration was mitigated in C3KO mice (Fig. 5E, G, I). Compared to NE WT mice, NE C3KO mice exhibited significantly larger MOC bouton densities in both apical and middle turns (Fig. 5F, G, I). These results suggest that C3 is required for noise-induced degeneration of MOC efferent boutons. However, despite the preservation of MOC terminal boutons, hearing impairments were not rescued in C3KO mice. This is consistent with prior works showing that the MOC system plays a modulatory role in auditory processing but does not directly impact absolute hearing thresholds (Guinan, 2017; Fuchs & Lauer, 2019; Mondul et al., 2024). In summary, C3 depletion selectively protects MOC efferent boutons from degeneration but does not prevent OHC loss or preserve hearing thresholds following severe NIHL. These findings suggest a targeted role for C3 in the efferent system that may be mechanistically distinct from its role in sensory hair cell pathology.

## Discussion

Previous studies have established that NE activates innate immune responses in the cochlea, including the upregulation of proinflammatory cytokines, infiltration of circulating immune cells, and activation of resident cochlear macrophages (Fujioka et al., 2006; Manickam et al., 2023, 2024; Pan et al., 2024; Rai et al., 2020; Wood & Zuo, 2017; Yang et al., 2015). However, the specific mechanisms that coordinate these immune responses, and their downstream effects on cochlear tissue remain unclear. Our findings of C3 activity following NE extend this framework by suggesting that the complement system may be the key mediator of tissue damage and clearance in the cochlea during NIHL. The complement system is a central component of innate immunity that orchestrates diverse immune functions, including damage recognition, opsonization, and inflammatory signaling (Dunkelberger & Song, 2010; Fishelson et al., 2001; Ricklin et al., 2016; Sarma & Ward, 2011; Vandendriessche et al., 2021). Although the expression of multiple complement genes was reported to be upregulated during both noise-induced and age-related hearing loss (Cai et al., 2014; Hoa et al., 2020; Su et al., 2020), the functional role of complement proteins—particularly C3—under these pathological conditions was not fully explored. Here, we show for the first time that C3 activation is tightly linked to noise pathology. Using a well-established NE model, we demonstrate that C3 activity is selectively and dynamically recruited to regions of damage in the cochlea. C3 accumulates in regions of OHC loss, opsonizes residual cellular debris and orphaned MOC efferent boutons, and participates in the process of IHC loss and ribbon synapse degeneration in an inducible model of selective cochlea damage. Notably, while genetic depletion of C3 did not prevent OHC loss or threshold shifts at 14 days after severe noise injury, it protected against degeneration of MOC boutons. These findings reveal a damage-dependent role for C3 in the injured cochlea and provide strong evidence that C3 plays an important role in coordinating damage recognition, debris clearance and cochlear immune responses, and contributing to post-injury tissue remodeling in the cochlea.

Versatile pattern recognition molecules of the complement system detect damage-associated molecular patterns and activate complement pathways, leading to the formation of C3 convertases (Dunkelberger & Song, 2010; Ricklin et al., 2016). While the native form of C3 is relatively inert, once activated and cleaved by C3 convertases following tissue injury, the resulting C3b rapidly opsonizes degenerating cells (Ricklin et al., 2016; Sarma & Ward, 2011). In our study, C3 accumulation was observed specifically in the basal cochlear turn, where OHC loss and synaptopathy were most severe, and was absent in the apical and middle turns unless damage was present (Fig. 1C-L). This damage-restricted pattern of C3 activation supports the role of complement in damage recognition and suggests that C3 activation in the cochlea is tightly linked to localized tissue degeneration rather than being a generalized response to NE. Particularly, C3 deposits were in the gaps left by missing OHCs and co-localized with degenerating OHCs and residual Myo7a-positive debris (Fig. 1I, K-M; Fig. 2N-Q; Fig. 3E-M), suggesting that C3 actively targets and opsonizes dying cells or remnants for clearance. This is consistent with known roles of C3 in opsonizing apoptotic or necrotic tissue in other peripheral systems (Dunkelberger & Song, 2010; Fishelson et al., 2001). We also found that C3 robustly colocalized with orphaned MOC efferent nerve boutons. These boutons were present within OHC-depleted regions and were specifically encapsulated by C3 (Fig. 2A-B, H-M).

Morphologically, these boutons displayed two distinct patterns that differed from normal controls: shrinkage (type 1) or swelling and dispersion (type 2) (Fig. 2C-G), similar to synaptic changes reported in prior noise and age-related hearing loss models (Boero et al., 2018; Canlon et al., 1999; Fu et al., 2010). It suggests that C3 may act across multiple stages of MOC degeneration and mediate distinct degenerative pathways.

Despite being responsive to injury, the organ of Corti lacks resident immune cells and immune infiltration under physiological conditions (Hirose et al., 2005, 2017; Okano et al., 2008; Yang et al., 2015). While macrophages can’t infiltrate into the sensory epithelium, it remains unclear whether their processes extend into this region and directly interact with degenerating tissues.

Prior studies suggest that supporting cells, particularly Deiters’ cells (DCs), respond to acoustic trauma and serve as ‘amateur’ phagocytes that engulf and clear OHC debris (Abrashkin et al., 2006; Anttonen et al., 2014; Hirose et al., 2017; Monzack et al., 2015). However, the molecular mechanisms that regulate debris recognition and phagocytosis in this context remain poorly understood. Our observation of C3-opsonized structures within the organ of Corti suggests that C3 may mediate the clearance of damaged cells via interaction with C3 receptors such as CR3 (CD11b), which facilitates complement-dependent phagocytosis (Vandendriessche et al., 2021). CR3 is predominantly expressed on immune cells, particularly macrophages, and is rarely reported on non-immune cells (ROSS & VĚTVIČKA, 1993; Vandendriessche et al., 2021). Its expression in the cochlea has not been well characterized. Future studies should investigate the expression and function of complement receptors on DCs and other immune-like cells in the cochlea to determine whether these cells engage with C3-opsonized debris following noise trauma.

Previous studies have described an overall timeline of immune responses in the cochlea following NE, with cytokine and chemokine expression peaking at 1–2 days and again around 7 days post-exposure, with immune cell infiltration peaking at 4–5 days (Kaur et al., 2015; Tan et al., 2016; Wood & Zuo, 2017). Remarkably, we observed C3 activation as early as D1 opsonizing damaged OHCs, albeit low in number, followed by sustained pattern of C3 activation peaking at D14 and persisting through D60 (Fig. 3C, E, K). The peak activation coincided with the observed period of maximal OHC degeneration (Fig. 3B-C), highlighting the damage-dependent activity of C3. Unlike previous studies (Hu et al., 2002; Ping Yang et al., 2004), we observed less OHC loss at early stages (D0-D5) after NE (Fig. 3B, D, E, H), which was likely due to the lower NE level and different frequency range in our experiment. Over time, C3 was first found to opsonize degenerating OHCs at D1 (Fig. 3E-G), shifted to OHC debris at D5 (Fig. 3H-J) and continued to D14 and D60 (Fig. 2N; Fig. 3K-M). C3 opsonization of MOC efferent boutons only became prominent at D14, with a time delay after OHC damage (Fig. 2A-B, H-M; Fig. 3F, J), suggesting distinct temporal windows of complement activity that align with the staged progression of cochlear pathology following NE. Previous studies reported that activated macrophages persist in the cochlea for at least 20 days post-injury, maintaining a state of chronic inflammation (Rai et al., 2020; Zhang et al., 2020). Our findings of persistent C3 activation at D60, well beyond the typical resolution phase of inflammation, raise the possibility that the complement system contributes to this chronic, low-grade immune activity in the cochlea. The prolonged presence of C3 may also reflect limited capacity for debris clearance in the cochlea, possibly due to the lack of professional phagocytes and the reliance on supporting cells for debris removal (Abrashkin et al., 2006; Anttonen et al., 2014; Hirose et al., 2017; Monzack et al., 2015).

Noise insult is known to damage other structures of the sensory epithelium including IHCs and cochlear synapses. However, much of the C3 activity we observed was in the OHC region, likely due to the lack of IHC damage and the instantaneous loss of cochlear synapses in our experiments that prevented the observation of C3 activity on both structures. Using the DT-mediated IHC ablation model, we found that C3 was recruited to IHCs undergoing cell death (Fig. 4D-E). Moreover, C3 colocalized with degenerating ribbon synapse components, including orphan presynaptic CtBP2 and postsynaptic Homer1 puncta (Fig. 4F-L). These findings suggest that C3 does activate promptly upon damages of IHCs and cochlear synapses, and may mediate synapse pruning in the cochlea, which was observed in non-auditory central nervous systems (Schafer et al., 2012; Stevens et al., 2007).

To further investigate C3 function in the cochlea, we used C3-deficient mice to test whether C3 contributes to hearing loss or tissue degeneration. These mice have C57BL/6J genetic but exhibit a similar pattern of cochlear pathology and auditory impairment (Wu et al., 2024). Consistent with our findings in CBA/CaJ mice, we observed prominent C3 accumulation in the cochleae of noise-exposed WT C57BL/6J mice (Fig. 5D, F, H). In contrast, C3-deficient mice showed complete absence of C3 upregulation (Fig. 5G-H), confirming the validity of our mouse model. Surprisingly, C3-deficient mice exhibited normal ABR thresholds and OHC counts under normal condition (Fig. 5A-B), suggesting that C3 may not be essential for cochlear development or normal function. A previous study reported that complement factor B (fB)-deficient mice develop hearing loss, with structural abnormalities in the auditory nerve and cochlear lateral wall (Brown et al., 2023). Since fB is a key component of the alternative complement pathway and facilitates the formation of C3 convertases (Ricklin et al., 2016; Sarma & Ward, 2011), this finding underscores the importance of complement signaling in auditory function. However, the long-term consequences of C3 deficiency on healthy and injured cochlea remain to be fully determined.

Following NE, although both WT and C3KO mice exhibited comparable ABR threshold shifts and OHC loss (Fig. 5A-C, F-G), we observed a striking preservation of MOC bouton area in C3- deficient mice compared to WT mice (Fig. 5F-G, I), suggesting that C3 is required for the degeneration of MOC efferent terminals. These results imply that complement-mediated efferent synaptic loss occurs independently of hair cell degeneration and ABR threshold elevation. This form of MOC degeneration is in line with previous findings that the MOC system modulates auditory functions, but its disruption does not affect baseline hearing thresholds (Mondul et al., 2024). In the central nerve system, ablating C3 prevents age-related hippocampal neuronal decline (Shi et al., 2015), suggesting that C3 may be detrimental to cellular pathology. However, our NE protocol was designed to induce severe and irreversible cochlear damage (Fig. 5A-B), which might have masked more subtle protective effects of C3 depletion on OHC and ribbon synapse survival or hearing function. Future research using less noise injury is needed to determine C3’s effect in NIHL. Notably, we also found that MOC boutons degenerate in apical and middle cochlear regions where OHCs were preserved (Fig. 5F-G, I), indicating that MOC terminals may be independently vulnerable to acoustic trauma from OHC loss. This aligns with prior observation in noise-exposed guinea pigs, where reduced synaptophysin immunolabeling was detected despite intact OHCs (Niu et al., 2007). The precise mechanisms underlying efferent synapse loss and the factors influencing their degeneration remain unclear.

In summary, our results demonstrate that C3 plays a critical role in noise-induced cochlear pathology. C3 activity is spatially restricted to regions of tissue damage, temporally aligned with cell degeneration, and functionally required for MOC bouton loss. These findings highlight the importance of innate immune mechanisms in shaping cochlear outcomes after acoustic trauma and suggest that modulation of complement signaling may offer new strategies for preserving synaptic integrity and mitigating cochlear damage in sensorineural hearing loss.

## Author contributions

BS, ZG and RX designed the research. ZG and SL performed experiments. ZG, MS, AW and KG analyzed the data. ZG and RX drafted the paper. All authors edited and approved the final version of the paper submitted for publication. All authors qualify for authorship and agree to be accountable for the work.

## Acknowledgement

All confocal images were acquired using microscopes at the Campus Microscopy and Imaging Facility (CMIF) at The Ohio State University.

